# Gata2a maintains *cebpa* and *npm1a* in haematopoietic stem cells to sustain lineage differentiation and genome stability

**DOI:** 10.1101/2021.07.19.452890

**Authors:** Christopher B. Mahony, Boris Noyvert, Pavle Vrljicak, Sascha Ott, Martin Higgs, Rui Monteiro

## Abstract

The transcription factor Gata2 is required to produce and maintain haematopoietic stem and progenitor cells (HSPCs) in development and adult haematopoiesis. Mutations in GATA2 lead to GATA2 deficiency syndrome and predispose patients to acquire leukaemia. Here we use zebrafish gata2a enhancer deletion mutants and single cell transcriptomics to understand how GATA2 mediates survival and differentiation of haematopoietic stem cells in GATA2 deficiency. *Gata2a* mutants show marrow failure, neutropenia, B-lymphopenia and erythrocytosis from 6 months post-fertilization (mpf). Single cell transcriptional profiling of the adult kidney marrow demonstrated that HSPCs express elevated expression of erythroid- and decreased expression of myeloid genes, including *cebpa*. This is associated with a lineage skewing towards the erythroid fate at the expense of the myeloid fate. Thus, Gata2a is required to initiate and maintain lineage priming in HSPCs, favouring myeloid differentiation. Gata2a regulates expression of multiple targets associated with replication and DNA damage repair (DDR), including *npm1a*, a zebrafish NPM1 orthologue. Accordingly, mutant marrow cells show increased DNA damage associated with progressive loss of *npm1a* expression with age. This effect was replicated by inhibiting NPM1 activity in murine HPC7 progenitor cells. We propose that the impaired DDR leads to marrow failure in GATA2 deficiency. This leads to increased genomic instability in the surviving HSPCs, favouring acquisition of secondary leukaemogenic mutations.

## Introduction

The specification of haematopoietic stem and progenitor cells (HSPCs) from aortic endothelium during embryogenesis is a highly conserved process across many species. HSPCs mature and eventually seed their adult niche, the bone marrow in mammals and kidney marrow in zebrafish where they are able to self-renew, differentiate to any blood lineage and sustain haematopoiesis throughout life (Crane et al., 2017; Sawai et al., 2016). Lineage commitment and differentiation is tightly controlled by key genes and transcription factors.

GATA2 is a key haematopoietic transcription factor and is required for the formation and lineage output HSPCs and myelopoiesis (de Pater et al., 2013; Gao et al., 2013; Rodrigues et al., 2008). Loss of GATA2 expression due to GATA2 haploinsufficiency mutations in coding or enhancer regions causes haematopoietic disorders collectively referred to as GATA2 deficiency syndromes (Dickinson et al., 2014; Spinner et al., 2014). These patients are predisposed to mycobacterial or viral infections, suffer from lymphedema and develop marrow failure (McReynolds et al., 2018). The majority of GATA2 deficiency patients with inherited germline GATA2 mutations (~75%) develop early onset myelodisplastic syndrome (MDS) and Acute Myeloid Leukaemia (AML) at a median age of 20 (Hirabayashi et al., 2017; Wlodarski et al., 2017). Existing mouse models fail to mimic the disease progression observed in humans; they either show no haploinsufficiency phenotype, are embryonic lethal or require exogenous stress to reveal haematopoietic repopulation defects (Gao et al., 2013; Johnson et al., 2012; Soukup et al., 2019). In contrast, by deleting a conserved intronic endothelial enhancer in the zebrafish *gata2a* locus (*gata2a*^-/-^ mutants hereafter), we generated a model of *GATA2* deficiency (Dobrzycki et al., 2020) that show marrow hypocellularity, neutropenia and propensity to infections and to develop an AML-like phenotype in the adult kidney marrow (Dobrzycki et al., 2020). Here we make use of this model to investigate disease onset and study the transcriptional changes that occur at the single cell level in mutant haematopoietic cells to understand how *gata2a* deficiency leads to perturbed haematopoiesis, marrow failure and predisposition to MDS/AML.

We demonstrate that *gata2a*^-/-^ mutants show marrow hypocellularity from 6 months post-fertilization (mpf), accompanied by neutropenia, monocytopenia and increased numbers of erythrocytes. Single cell transcriptional profiling of whole kidney marrow (WKM, equivalent to bone marrow in humans) enabled us to identify all the major haematopoietic populations in the WKM and demonstrated that the neutropenia and erythrocytosis phenotypes are detectable at a molecular level at 5mpf, prior to the marrow failure and cytopenia phenotypes. *Gata2a*^-/-^ mutant HSPCs are mis-programmed, expressing increased levels of erythroid lineage genes (*hemgn, igfbp2a, hbba1, hbaa1*) and decreased expression of myeloid genes, including *lcp1, npm1a* and *cebpa*. Pseudotime analysis showed that loss of *cebpa* in *gata2a*^-/-^ HSPCs correlated with increased expression of erythroid lineage genes, suggesting that *cebpa* is a downstream effector of *gata2a* in HSPCs. In addition, mutant HSPCs showed decreased expression of key replication and DNA damage repair genes, including *npm1a*, indicating a possible mechanism for DNA damage repair in the bone marrow failure phenotype in GATA2 deficiency patients. Indeed, we show that *npm1a* expression is lost as disease progresses and correlates with the appearance of DNA damage in WKM and murine HPC7 multipotent progenitor cells. Taken together, *gata2a* is required to initiate and maintain lineage priming in HSPCs through *cebpa*, favouring the myeloid fate over the erythroid or B-lymphoid fates. Furthermore, we propose that *gata2a* regulates HSPC survival and protects against bone marrow failure by regulating expression of essential DNA damage repair genes. In the absence of *gata2a*, the impaired DNA damage repair machinery leads to increased DNA damage and decreased HSPC survival.

## Results

### Gata2a directs myeloid lineage differentiation and maintains homeostasis in the WKM

Our previous study of zebrafish *gata2a* revealed that *gata2a* enhancer (i4 enhancer) mutants show impaired haemogenic endothelium (HE) programming but HSPC numbers in the embryonic niche recover by 5dpf (Dobrzycki et al., 2020). However, at 6 months post-fertilization (mpf), adults exhibit a hypocellular WKM, neutropenia, and by 9mpf 1 in 10 show >98% blasts in the WKM, a hallmark of AML that indicated that the enhancer deletion mutants displayed key characteristics of GATA2 deficiency in humans (Dobrzycki et al., 2020). To uncover the mechanism underlying disease progression in GATA2 deficiency, we first determined the onset of the hypocellularity phenotype by analysing WKM from 4, 5 and 6mpf *gata2a* mutants by WKM smears and cell counts (Fig. 1A,B). These experiments revealed that the marrow hypocellularity only became apparent by 6mpf (Fig. 1 A,B). However, the hypocellular phenotype was detectable in *gata2a*^+/-^ heterozygous and was indistinguishable from homozygous mutants, indicating that maintenance of normal WKM cell numbers is very sensitive to the dose of *gata2a* (Fig. 1A, B). To examine how HSPC differentiation was affected, *gata2a*^-/-^ mutants were crossed to transgenic lines to examine haematopoietic populations in the WKM as previously established (Traver et al., 2003). We looked at neutrophils (*mpx:GFP*, myeloid gate) (Renshaw et al., 2006), macrophages (*mpeg1.1:GFP*, myeloid gate), B cell (*mpeg1.1:GFP*, lymphoid gate) (Ferrero et al., 2020) and erythrocytes (*gata1:DsRed*, erythroid gate) (Yaqoob et al., 2009) at 6mpf (Fig. 1C). As expected, we found decreased numbers of *mpx:GFP^+^* neutrophils in *gata2a*^-/-^ (Fig. 1D). In addition, we found decreased *mpeg1.1:GFP*^+^ B cells in the lymphoid gate (Fig. 1F) and increased *gata1a:DsRed^+^* cells in the erythroid gate (Fig. 1G). Together, these experiments showed that Gata2a is required to maintain haematopoietic cell numbers in the WKM and for correct lineage differentiation in the adult marrow. No significant skewing in lineage differentiation was found in *gata2a* heterozygous mutants, indicating that maintaining cellular homeostasis in the WKM requires higher levels of Gata2a and is more sensitive to Gata2a dosage than lineage determination.

**Figure 1.**
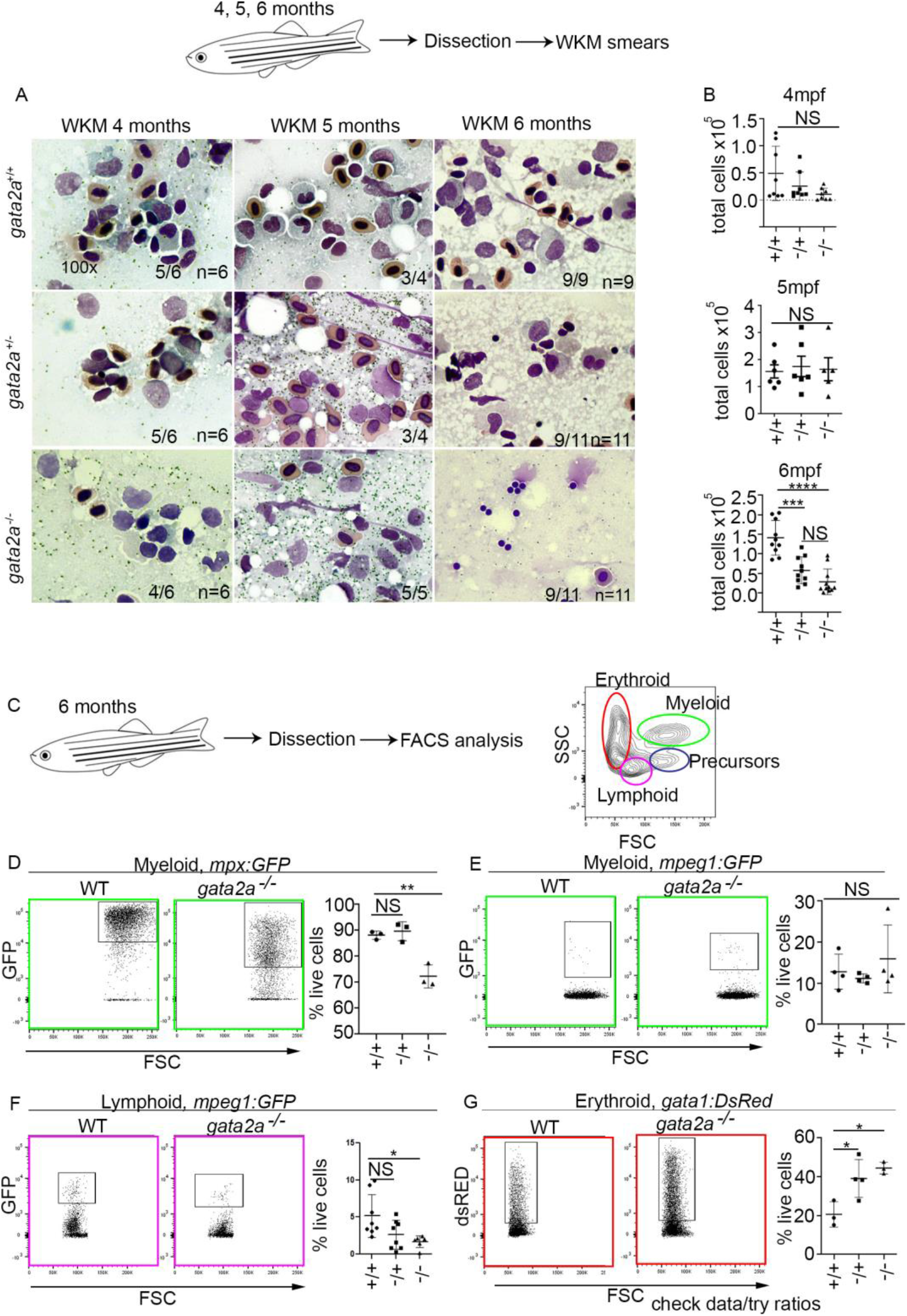
Loss of *gata2a* results in hypocellular marrow at 6mpf. (A) WKM smears and May Grunewald/Giemsa sating at 4, 5 and 6mpf. (B) total WKM cell counts at 4, 5 and 6mpf. (C) Flow cytometry gating strategy. (D-G) Analysis of different lineages using either *mpx:GFP, mpeg1:GFP* or *gata1:DsRed* transgenic lines. D, gata2a^+/+^ vs gata2a^+/-^: p=0.82, gata2a^+/+^ vs gata2a^-/-^: p=0.0024. E, gata2a^+/+^ vs gata2a^+/-^: p=0.87, gata2a^+/+^ vs gata2a^-/-^: p=0.64. F, gata2a^+/+^ vs gata2a^+/-^: p=0.28, gata2a^+/+^ vs gata2a^-/-^: p=0.0239. G, gata2a^+/+^ vs gata2a^+/-^: p=0.025, gata2a^+/+^ vs gata2a^-/-^: p=0.01. Analysis was completed using one-way ANOVA, NS: P>0.05, *: P<0.05.

### Single cell RNAseq analysis revealed lineage skewing towards the erythroid fate at the expense of myeloid cell fates

To investigate how *gata2a* regulates WKM cellularity and lineage differentiation, we undertook transcriptional profiling of the WKM at single cell resolution at 5mpf, prior to the onset of the detectable lineage skewing and hypocellularity phenotypes by flow cytometry. We took two adult fish per genotype, dissected and pooled the WKM and sorted live cells for scRNAseq using the 10X Chromium platform. Analysis estimated 3197 WT cells and 5838 gata2a^-/-^ cells. We could readily identify cell clusters representing all the major haematopoietic and kidney cell populations present in the WKM (Fig. 2A), consistent with recent studies (Moore et al., 2016; Tang et al., 2017). We defined ‘core HSPCs’ (cHSPCs) by expression of *myb, lmo2, cebpa, meis1b, gata2b* and *runx1* (Fig. S1A, 2B). They were also enriched in other known embryonic HSPC markers such as *gfi1aa, dnmt3ba, angpt1*, and *si:dkey-261h17.1* (identified as the zebrafish orthologue of *CD34*, zfin.org, hereafter referred to as *cd34-like*, or *cd34l*) (Fig. S1A, 2B). Phylogeny and synteny analyses revealed the conservation of the CD34 antigen as defined by ENSEMBL (Fig. S2A-C). We also defined a second HSPC population as ‘myeloid-primed’ HSPCs (mHSPCs) based on their lack of expression of pluripotency markers (such as *gata2b, runx1*, and *meis1b*) and their retention of myeloid-biased transcription factors (*lmo2, cebpa* and *myb*) and higher expression levels of *c1qa* and *mafba* (Fig. S1A). We then defined the remaining populations based on their expression of key marker genes (*igfbp1a, hemgn, klf1:* Erythroid progenitors (Ery-P), *c1qa, mfap4, mafba:*monocytes/macrophages, *mpx, mmp13a, vamp8*: neutrophils, *nkl*.3, *il2rb*: NK/Tcells, *pax5, cd37, cd79b:* B cells, *sox8b, tekt1*: multi-ciliated kidney cells, *spink2.2, viml:* kidney tubule cells) (Fig. 2B, S1A). When we compared wildtype and gata2a^-/-^ WKM we found a striking increase in erythroid cells and a decrease in neutrophils (Fig. 2C), indicating that the lineage skewing observed at 6mpf (Fig. 1) could be detected at the cellular and molecular level at 5mpf (Fig. 2) by single cell RNAseq.

**Figure 2.**
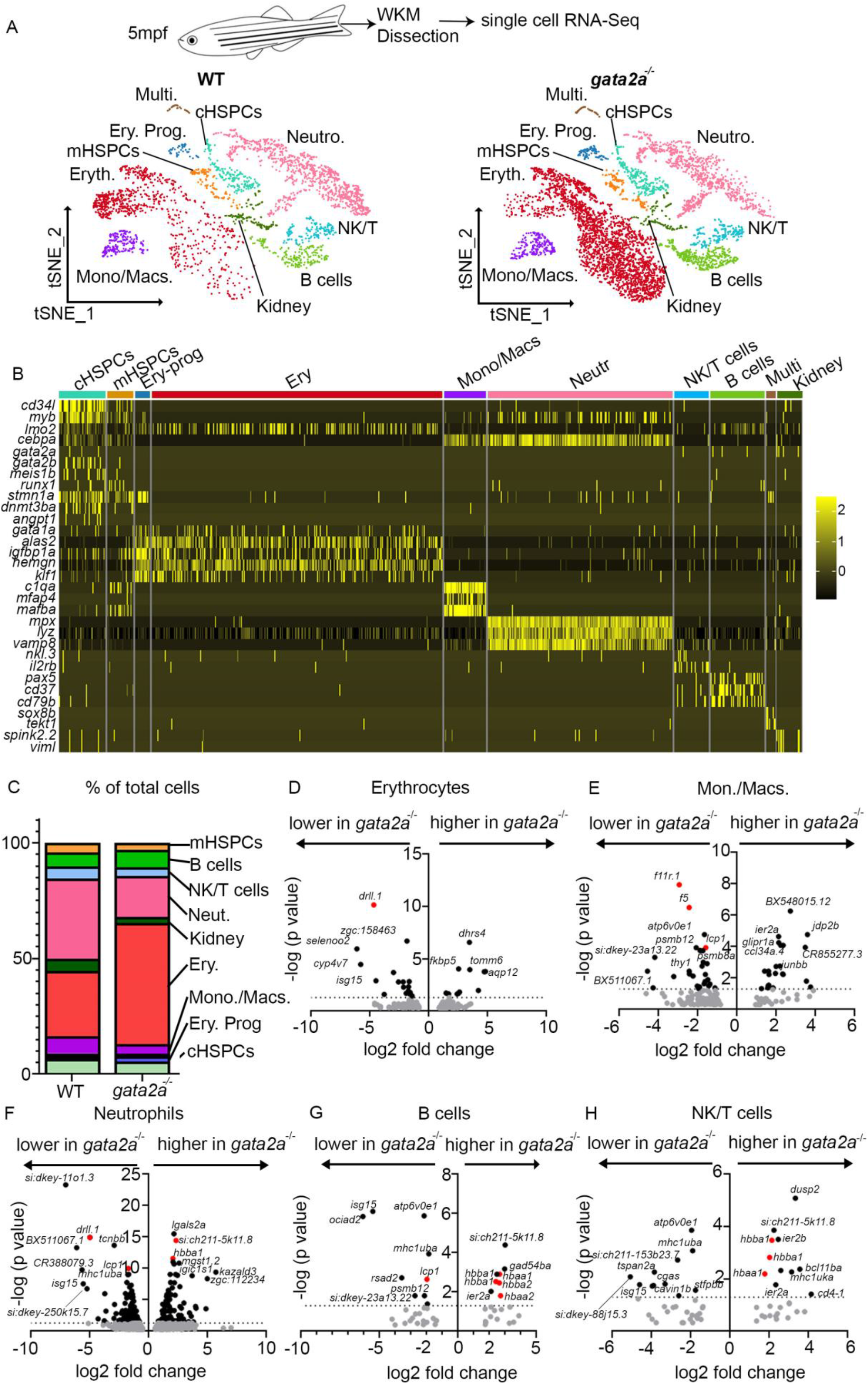
*gata2a* is required for lineage specification and transcriptional regulation in WKM haematopoietic populations. (A) scRNA-Seq and tSNE clustering of WKM haematopoietic populations. (B) Mean (Log2) expression levels of indicated genes in specific populations. cHSPCs: core HSPCs, mHSPCs: myeloid primed HSPCs, Ery Prog: erythroid progenitors, Mono/Macs.: Monocytes/Macrophages, Eryth: erythroid cells, Neut.: Neutrophils, NK/T: NK/T cells, Multi.: Multi ciliated cells. (C) Percentage of WKM populations of total WT or *gata2a*^-/-^ cells. (D-H) Volcano plots of log2 fold change vs –log10 (p value, determined by using the Benjamini-Hochberg correction for multiple tests) in different WKM populations. Each dot represent a gene. Dotted line indicates p=0.05 with grey genes falling below this threshold.

When then examined differential gene expression in neutrophils, monocytes/macrophages, erythrocytes, NK/Tcells and B cells. Within the Ery-P we found no differentially expressed genes. Surprisingly, despite their increase in number, relatively few genes were significantly changed in erythrocytes (Fig 2D). They showed a decrease in *drll*.1, a transcription factor required for erythropoiesis (Kobayashi et al., 2020) and ectopic expression of *aqp12* and *dhrs4*, suggesting that although their number was increased, their function was likely impaired (Fig. 2D). Monocytes showed a decrease in the myeloid markers *lcp1, f5* and *f11r.1* (AKA *jam-a*) (Luissint et al., 2019) in *gata2a*^-/-^ cells indicating altered differentiation and function (Fig. 2D). *gata2a*^-/-^ mutant neutrophils also expressed less *lcp1* and *f5* and more *si:ch211-5k11.8* (Orthologous to human HBZ-haemoglobin zeta, zfin.org) and *hbba1* (Fig. 2E). Mutant B cells expressed less *lcp1* and more *hbba1, hbaa1, hbba2, hbaa2* and *si:ch211-5k11.8* (Fig. 2F). Mutant NK/T cells expressed more *hbba1, hbaa1* and *si:ch211-5k11.8* (Fig. 2E). In summary, the myeloid and lymphoid haematopoietic populations show downregulation of genes of myeloid lineage and ectopic upregulation of genes of the erythroid lineage. Taken together, this suggested that Gata2a, mainly expressed in the cHSPC population (Fig. 2A, S1B), is required to maintain transcriptional states of myeloid and erythroid genes in the most differentiated haematopoietic populations and that aberrant gene expression in differentiated blood lineages could be traced back to the HSPC populations. To investigate this, we next examined differential gene expression in cHSPCs and mHSPCs (Fig. 3). Mutant cHSPCs showed a decrease in *drll.1*, as well as *atp6V0e1* and *mhc1uba*, and an increase in the HBZ orthologue, *si:ch211-5k11.8* (Fig. 3A). GO term enrichment analysis highlighted only very few pathways, suggesting that cHSPCs remain relatively unaffected (Fig.3C, D). Many more genes were differentially expressed in mHSPCs (Fig. 3B). We found that globin genes (*e.g.hbba1, hbaa1, hbba2, hbaa2*: Figure 3G), as well as known erythroid regulators like *hemgn* (Peters et al., 2018), *igfbp1a, znlf2a* and *rhag*, were all upregulated in mHSPCs, which was confirmed by GO term enrichment analysis which highlighted that upregulated genes were enriched in pathways for erythrocyte differentiation (Fig. 3E). By contrast, *cebpa* (Figure 3G) and myeloid lineage markers like *lcp1, lsp1b* and *f5* (Fig. 3B) were downregulated. *cebpa* is a key regulator of granulocyte and monocyte differentiation and primes HSPCs towards the myeloid fate (Avellino and Delwel, 2017; Dai et al., 2016; Hasemann et al., 2014; Suh et al., 2006). Thus, the differentiation skewing towards the erythroid lineage at the expense of myeloid lineages in *gata2a*^-/-^ mutants is likely driven by the loss of *cebpa* in mHSPCs (Fig. 3B). We found that critical regulators of DNA replication (e.g. *mcm3, mcm5*: Fig. 3G) were downregulated in mHSPCs, suggesting a decrease in proliferation (Shinya et al., 2014; Snyder et al., 2005) consistent with the cytopenic phenotype noted at 6mpf. In addition we noted a decrease in genes required for DNA repair and genome stability, such as *hells* (Kollárovič et al., 2020), *npm1a* (Grisendi et al., 2005; Koike et al., 2010) and *ncl* (Kawamura et al., 2019; Scott and Oeffinger, 2016) (Fig. 3G). This was confirmed by GO term enrichment analysis which highlighted that downregulated genes were enriched for pathways in DNA replication initiation, chromatin assembly/disassembly and methylation (Fig. 3F). Other GO terms included ribosome biogenesis, highlighted due to the previous role of *NPM1* in ribosome biogenesis (Lindström, 2011), centromere formation and histone chaperoning (Grisendi et al., 2005; Okuwaki et al., 2001).

**Figure 3.**
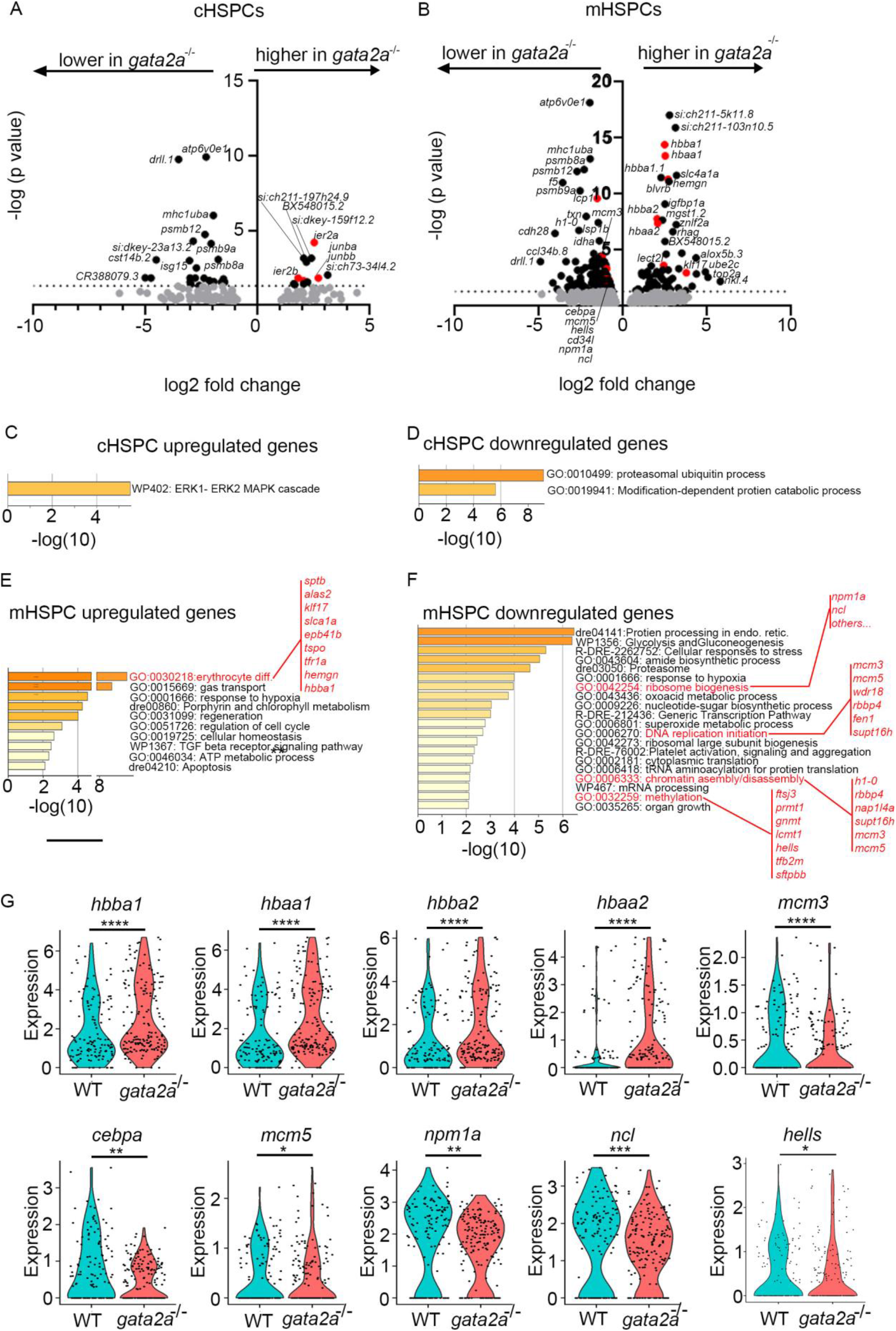
Loss of *gata2* in mHSPCs dysregulates the expression of differentiation and DNA repair genes. (A, B) Volcano plots of log2 fold change vs –log10(p value, determined by using the Benjamini-Hochberg correction for multiple tests) in cHSPCs (core HSPCs) and mHSPCs (Myeloid primed HSPCs) populations. Each dot represent a gene. Dotted line indicates p=0.05 with grey genes falling below this threshold. (C-F) GO term enrichment (performed using metascape.org) analysis of up or down regulated genes in either cHSPCs or mHSPCs with a p value < 0.05. (G) Expression levels were calculates using LogNormalize (Seurat: R package), by dividing feature counts by the total counts for that cell and then multiplied by the scale factor. This is then natural-log transformed using log1p. ****: p<0.0001, ***:p<0.001 **:p<0.01, *:p<0.05 determined by using the Benjamini-Hochberg correction for multiple tests.

We conclude that *gata2a* expression in cHSPCs is required to establish a transcriptional network that controls erythroid and myeloid fate through *cebpa*, HSPC fitness and survival via regulation of metabolic pathways and the expression of DNA repair genes.

### Pseudotime analysis shows that lineage skewing in gata2a^-/-^ originates from HSPCs and correlates with loss of *cebpa* expression

We noted that a number of genes were consistently up-(e.g. *ier2a*, *junbb*) or downregulated (*e.g. atp6V0e1, mhc1uba*) across several haematopoietic populations including HSPCs (Fig. S2, Fig. 2, Fig. 3). To understand the dynamics of gene expression changes found in *gata2a*^-/-^ mutants, we examined gene expression during HSPC differentiation using pseudotime analysis with Monocle 3 (Cao et al., 2019; Qiu et al., 2017a; Qiu et al., 2017b; Trapnell et al., 2014). For that, we first plotted the WKM populations using UMAP clustering and identified HSPCs based on their expression of *lmo2, myb, gata2b, cd34l, meis1b* and *cebpa* (Fig. S3A, B). We repeated the UMAP clustering in Monocle 3 and determined the expression of the same genes (Fig. S3C), which allowed us to select a broad HSPCs population (Fig. S3D) for further pseudotime analysis. We then reduced dimension and defined five distinct HSPC populations based on marker expression (Fig. 4A). We first defined cHSPCs based on their higher expression of *cd34l, meis1b, gata2b* and *lmo2* (Figure S4A). mHSPCs were defined by higher expression of *cebpa, myb*, and *c1qbp* along with more restricted *spi1b* expression (Fig. S4B). Proliferating HSPCs were defined by high expression of histone genes (Fig. S4C). One population was enriched in haemoglobins which we classified as erythrocyte biased HSPCs (eryHSPCs, Fig. S4D). A fifth, small population of HSPCs was enriched in lymphoid genes (*rag1, rag2, cxcr4b, btg1*) and thus defined as lymphoid-biased HSPCs (lymphHSPCs, Fig. S4E). We then examined expression of key myeloid and erythroid genes in pseudotime by classifying cHSPCs as the differentiation origin. For this analysis, we excluded proliferating HSPCs. To understand how the expression of myeloid and erythroid genes changes with differentiation, we visualised the expression of *cebpa*, *hbba2*, *hbaa2* and *hemgn* in pseudotime. We found that wild type cHSPCs and mHSPCs express *cebpa* at similar levels whereas it is downregulated as they differentiate towards eryHSPCs (Fig. 3B). By contrast, *gata2a*^-/-^ cHSPCs and mHSPCs show severely decreased *cebpa* expression that is downregulated early on in pseudotime. The erythroid genes *hbba2, hbaa2* and *hemgn* are expressed at low levels in wildtype cHSPCs and mHSPCs and are upregulated as HSPC differentiate to eryHSPCs. Both *hbba2* and *hbaa2* are upregulated in *gata2a*^-/-^ cHSPCs and robustly upregulated in mHSPCs and eryHSPCs (Fig. 3B). These results show that loss of *cebpa* as HSPCs differentiate correlated with enhanced erythroid marker expression and reduced myeloid marker expression, suggesting that a Gata2a-Cebpa axis mediates myeloid lineage priming in HSPCs.

**Figure 4.**
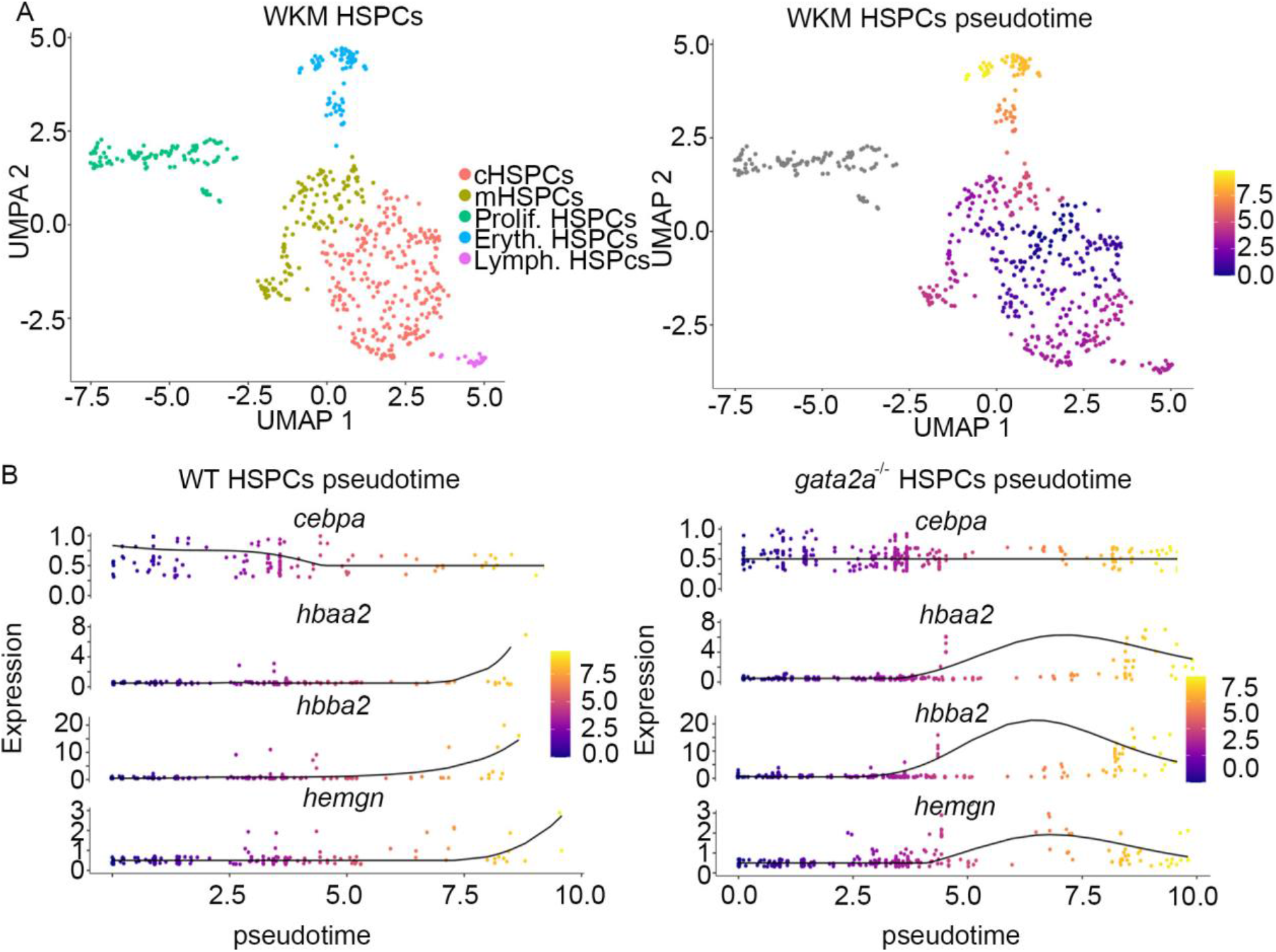
*gata2a* regulates *cebpa* expression as HSPCs differentiate. (A) UMAP plots of selected HSPCs clustered into different sub populations and ordered in pseudotime with cHSPCs defines as the point of differentiation origin. cHSPCs: core HSPCs, mHSPCs: myeloid primed HSPCs, Prolif. HSPCs: proliferating HSPCs, Eryth. HSPCs: erythroid primed HSPCs, Lymph. HSPCs: lymphoid primed HSPCs. (B) WT and *gata2a*^-/-^ expression of *cebpa, hbaa2, hbba2* and *hemgn* in pseudotime.

### The nucleophosmin orthologue *npm1a* is highly enriched in HSPCs

Mutations in the C-terminal nuclear localization sequence of nucleophosmin (NPM1) are associated with myeloid and lymphoid malignancies (Sportoletti et al., 2008) and are found in around 30% of AML cases (Falini et al., 2005). Furthermore NPM1 is associated with maintenance of genome stability and required for DNA repair (Grisendi et al., 2005; Koike et al., 2010). At 5mpf, expression of the NPM1 orthologue, *npm1a* expression was most highly enriched in cHSPCs and mHSPCs, along with *cd34l* (Fig. S5A, A’). *npm1a* expression was decreased in *gata2a*^-/-^ mHSPCs (Fig. 3B,G) at 5mpf, suggesting that it may play a role in disease progression in GATA2 deficiency. To investigate this, we first asked whether *npm1a* expression was detectable in HSPCs isolated from WKM by expression of low levels of GFP driven by the CD41 promoter (CD41:GFP^low^ HSPCs) (Ma et al., 2011). Because our experiment was performed using wildtype and *gata2a*^-/-^ WKM on a *cd41:GFP* transgenic background, we mapped *gfp* expression and confirmed that it was highly enriched in mHSPCs and cHSPCs (Fig. S5A, A’). We grouped cHSPCs and mHSPCs into HSPCs and partitioned the resulting population into three groups: GFP^negative^ (expression less than 0.2 (log2 scale)), GFP^low^ (expression from 0.2 to 4 (log2 scale)) and GFP^high^ (expression > 4 (log2 scale)). We then identified the global markers associated with each population compared to the WKM. The GFP^negative^ HSPCs displayed enrichment in erythroid markers (*znfl2a, mycb, hbae1.3*) and *npm1a* (Fig. S5B). As expected, the GFP^low^ population was highly enriched in HSPC-associated markers, including *gata2b, cd34l, runx1t1* and *npm1a*. The GFP^high^ population expressed higher levels of thrombocyte-associated genes including *mpl* and *gp1bb* (Figure S5D, E) but lacked *npm1a* and *cd34l*. This confirmed that *npm1a* is expressed in the CD41:GFP^low^ HSPC population and allowed us to examine whether loss of *npm1a* correlates with disease progression.

We took two *cd41:GFP* and two *cd41:GFP;gata2a*^-/-^ adults at 7mpf, dissected WKM and sorted live, CD41:GFP^low^ cells from the lymphoid and progenitor gates (Fig. 5A). FACS analysis showed a decrease in the number of CD41:GFP^low^ cells in the mutant samples, as expected (Fig. 5A). We then processed the sorted cells for scRNA-Seq. Analysis estimated 700 WT cells (median UMI counts per cell=341, mean reads per cell= 123905, median genes per cell= 156) and 1117 gata2a^-/-^ cells (median UMI counts per cell=258, mean reads per cell= 75715, median genes per cell= 128). Due to low fraction reads, we chose cells with at least 1000 UMI counts in R and then exported extracted barcoded reads to be retained for further analysis in cell ranger/Loupe. We then analysed 106 wildtype cells (pooled from both samples) and 72 mutant cells (pooled from both samples) that revealed two distinctly separated GFP^low^ HSPC populations between wildtype and *gata2a*^-/-^ HSPCs (Fig. 5B). tSNE plot of the cells showed two distinct clusters made up exclusively of either WT or gata2a^-/-^ cells (Fig. 5C). Differential expression analysis showed a strong down regulation of *npm1a* in *gata2a*^-/-^ HSPCs (Fig. 5D, E) and upregulation of innate immune response genes (*lyz* and *lect2l*, Fig. 5D, F: a likely consequence of a shift to myeloid blast accumulation preceding MDS/AML onset). Consistent with the erythroid skewing found in 5mpf WKM, *hbba2* and *hbaa2* were still upregulated in *gata2a*^-/-^ HSPCs at 7mpf (Fig. 5D, F). Thus, we concluded that disease progression correlated with a decrease in *npm1a* expression in HSPCs.

**Figure 5.**
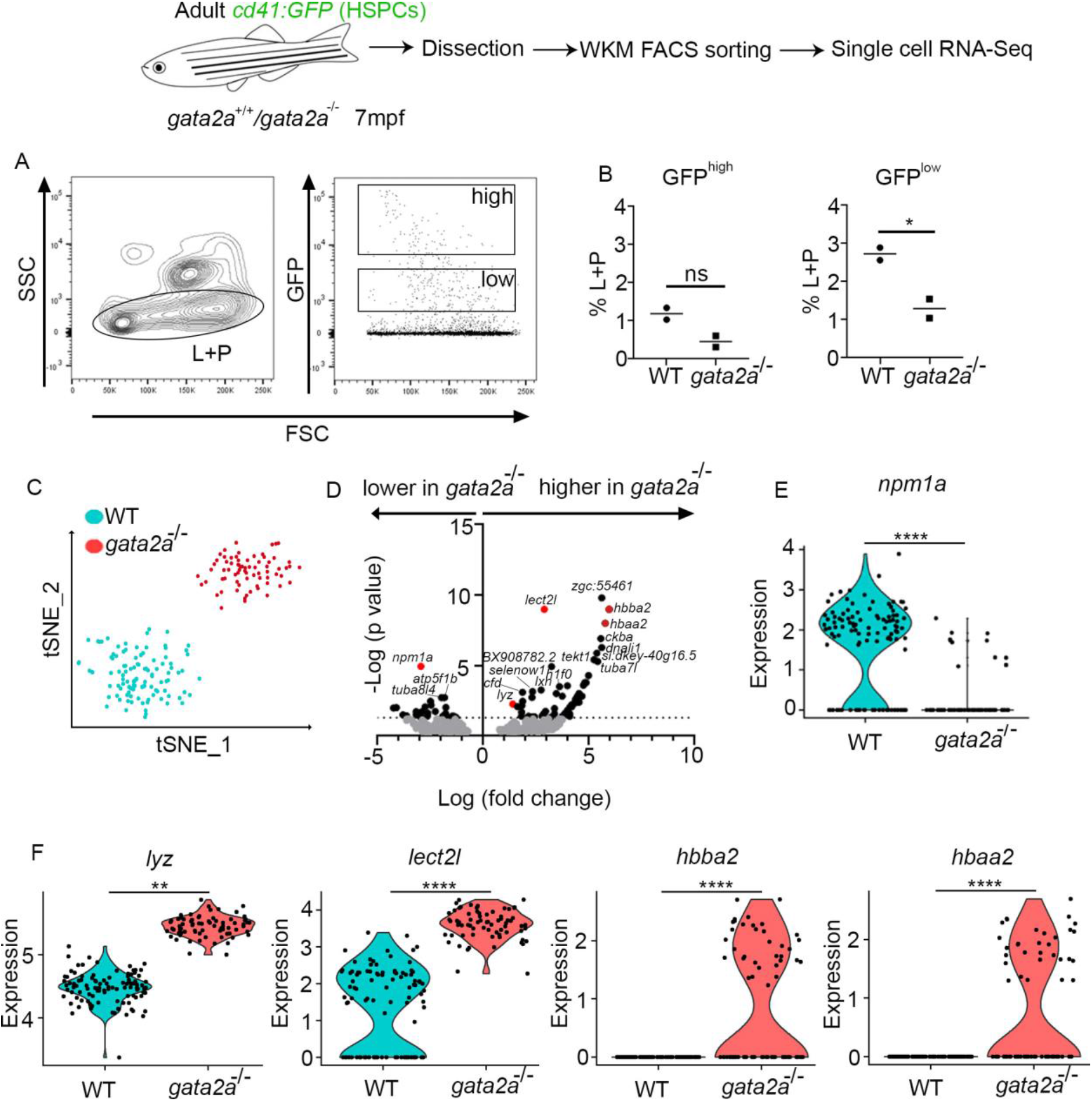
*npm1a* is down regulated as disease progresses in *gata2a*^-/-^ adults. (A) FACS gating strategy to analyse/isolate *cd41:GFP^low^* from 7mpf adults. L: lymphoid cells, P: progenitor cells. (B) analysis of GFP^high^ and GFP^low^ fraction. (C) scRNA-Seq and tSNE clustering of WT and *gata2a*^-/-^ cells. (D) Volcano plots of log2 fold change vs – log10(p value). (E) Violin plot (each dot represent a single cell) of *npm1a* expression in WT and *gata2a*^-/-^ cells. (F) Violin plot (each dot represent a single cell) of *lyz, lect2l, hbba2*, and *hbaa2* expression in WT and *gata2a*^-/-^ cells. Expression levels were calculates using LogNormalize (Seurat: R package), by use dividing feature counts by the total counts for that cell and then multiplied by the scale factor. This is then natural-log transformed using log1p. ****: p<0.0001, **:p<0.01, determined by using the Benjamini-Hochberg correction for multiple tests.

### Loss of *npm1a* correlates with DNA damage

Loss of Npm1 function in a murine hypomorphic mutant leads to genomic instability, impaired repair of DNA double strand break damage and promotes oncogenesis (Andrade et al., 2020; Grisendi et al., 2005). Thus, we wanted to examine if the decrease in *npm1a* in HSPCs correlates with increased DNA damage in the *gata2a*^-/-^ mutant WKM. For this, we examined γH2AX staining, a marker of DNA damage (Sharma et al., 2012), in WKM smears at 6 and 12mpf (Fig. 6). As a positive control, irradiation with 10Gy induced nuclear γH2AX staining in a freshly isolated wildtype WKM smear (Fig. 6A). Wildtype WKM cells at 6 and 12 mpf showed no evidence of γH2AX staining (Fig. 6B). By contrast, we could readily detect γH2AX^+^ cells in WKM from gata2a^-/-^ adults at 6 and 12mpf (Fig. 6B, S6A). Flow cytometry analysis at 12mpf showed that increased γH2AX was also detected in gata2a^-/-^ WKM suspensions suggesting that loss of *gata2a* leads to increased DNA damage in HSPCs (Fig. 6C). To independently confirm these results in a different model, we examined γH2AX following NPM1 inhibition in murine haematopoietic precursor HPC7 cells (Pinto do et al., 1998). Inhibition with 1μM of the NPM1 inhibitor for 24h showed an increase in γH2AX (Figure 6D), indicating that NPM1/npm1a is required to prevent increased DNA damage and maintain genomic stability in HSPCs.

**Figure 6.**
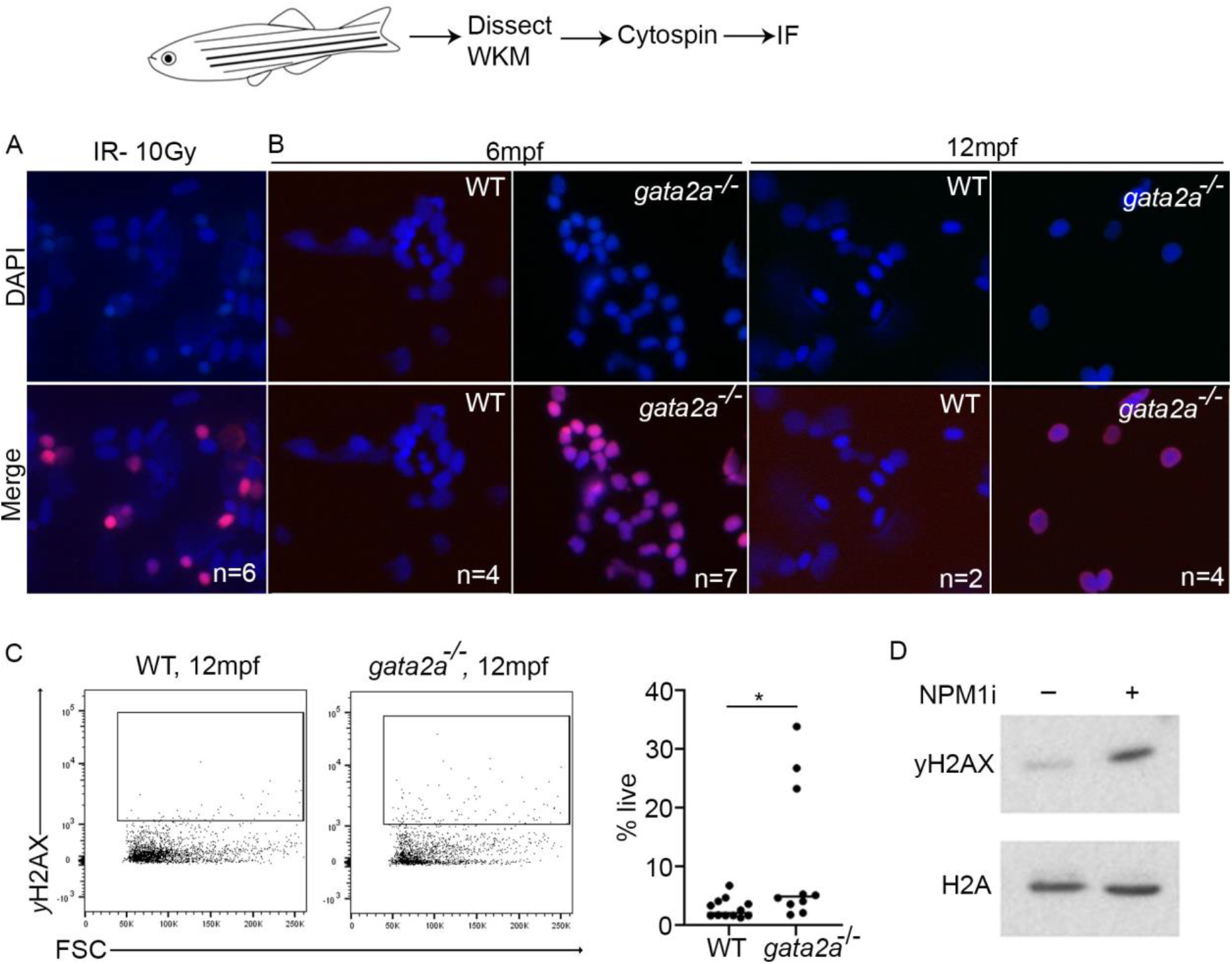
Loss of *npm1a* correlates with the accumulation of DNA damage. (A) Control irradiation performed at 10 Gy on WKM cells and immunofluorescence against DAPI and γH2AX (B) WKM from 6mpf and 12mpf WT or gata2a^-/-^ adults and immunofluorescence against DAPI and γH2AX. Imaging is at 40x magnification. (C) Flow cytometry analysis of γH2AX expression in 12mpf WKM suspension. *p*=0.0296. (D) Western blot of γH2AX and H2A expression in in HPC7 cells after 1μM NPM1 inhibitor for 24h. *:*p*<0.05, determined by using an unpaired, two tailed t-test.

In summary, we concluded that *gata2a* regulates *cebpa* and *npm1a* in HSPCs. Loss of *cebpa* results in skewed differentiation towards erythroid lineage at the expense of the myeloid lineages (neutrophils and monocytes), whereas loss of *npm1a* results in the inability to efficiently repair DNA damage in HSPCs. We propose that this results in increased genomic instability leading to the accumulation of somatic mutations and disease progression from a first phase of hypocellular marrow to a second phase of myeloid proliferation in HSPCs and onset of MDS/AML in GATA2 deficiency.

## Discussion

We have shown that deletion of a conserved *gata2a* enhancer (the i4 enhancer) leads to hypocellular marrow and a clear disease phenotype from 6mpf. Loss of the i4 enhancer leads to ectopic expression of erythrocyte genes in HSPCs and myeloid and lymphoid lineages and differentiation skewing from 5mpf as a result of decreased *cebpa* expression. Furthermore, decreased *npm1a* expression in mHSPCs correlates with the appearance of DNA damage in WKM cells and in murine haematopoietic progenitor cells, HPC7. These transcriptional changes lead to aberrant differentiation and defective DNA repair that ultimately trigger the marrow failure phenotype.

In our scRNA-Seq, we find less *cebpa* in *gata2a*^-/-^ mHSPCs, despite *gata2a* being mainly expressed in cHSPCs. Genetic and epigenetic analysis of CEBPa indicates that it acts as an HSC priming factor to promote myeloid differentiation and protects HSCs against apoptosis and maintains quiescence (Hasemann et al., 2014; Suh et al., 2006). We therefore hypothesise that *gata2a* sets up a transcriptional network in cHSPCs, possibly through chromatin remodelling, that is required for the expression of *cebpa* in mHSPCs. Furthermore, a downstream CEBPa enhancer is bound by GATA2 (Cooper et al., 2015), suggesting that the equivalent *cebpa* enhancer could be directly regulated by *gata2a* in zebrafish HSPCs. We therefore conclude that in our *gata2a*^-/-^ mutants, as cHSPCs begin to differentiate to mHSPC, lack the epigenetic cues that maintain *cebpa* expression and therefore differentiate to more erythroid lineages at the expense of myeloid ones (Suh et al., 2006). This is illustrated well by our pseudotime analysis of HSPC differentiation (Fig. 4) where *cebpa* expression is much lower in the *gata2a*^-/-^ mutants and erythroid genes are ectopically expressed.

Analysis of the *gata2a* zebrafish orthologue, *gata2b*, found that it was responsible for myeloid lineage commitment, over lymphoid and erythroid, when examining the transcriptional changes in immature *gata2b* deficient HSPCs (Gioacchino et al., 2021). Further analysis showed alterations in chromatin accessibility in early myeloid and lymphoid progenitors from whole marrow studies in a different *gata2b* mutant leading to defects in myeloid and lymphoid differentiation (Avagyan et al., 2021). As both *gata2a* and *gata2b* are expressed in HSPCs and that loss of either gene leads to a defect in myelopoiesis/lymphopoiesis we conclude that they have overlapping roles in HSPC biology and would function as “allelic paralogues” to recapitulate the human phenotype in germline heterozygous LOF mutations. However, our *gata2a* mutant (Dobrzycki et al., 2020) recapitulates a wider range of GATA2 dficiency syndrome symptoms including the recurrent infections, lymphedema, cytopenia and predisposition to MDS/AML and thus more faithfully recapitulates GATA2 deficiency syndrome.

In our characterisation of the WKM cell population by scRNA-Seq we identified the zebrafish orthologue of cd34 (*cd34l/ si:dkey-261h17.1*). Interestingly, the function of CD34 is to enable HSPCs to anchor to their marrow niche (Healy et al., 1995). We noted that cd34l was also modestly decreased in mHSPCs (Fig. 3B), which is likely to be contributing to the hypocellular phenotype seen in our model. We hypothesize that with less *cd34l*, HSPCs anchor themselves to the WKM niche less efficiently (Healy et al., 1995) and are no longer exposed to supportive niche signals and therefore differentiate and reduce their proliferation.

Appearance of MDS/AML in patients carrying germline GATA2 mutations is associated with secondary mutations in several genes (Hirabayashi et al., 2017; McReynolds et al., 2019; Wang et al., 2015; West et al., 2014). Loss of one Gata2 allele in a murine Cbfb-MYH11 fusion AML model leads to a ~2-fold increase in somatic mutations (Saida et al., 2020). While these studies provide an association between the genotype (mutated genes) and the phenotype (leukaemia), they fail to address how the identified mutations arise or how they affect gene expression to initiate or exacerbate disease progression.

Here, we have found that loss of *gata2a* leads to less *npm1a* and the acquisition of DNA damage. Although it remains unknown whether the regulation of *npm1a* by *gata2a* is direct, previous studies have shown that *NPM1* maintains genome integrity during cell cycle, inhibits apoptosis/tumour suppression through p53 and its loss of function is associated with myeloid and lymphoid malignancies (Colombo et al., 2002; Okuwaki, 2008; Sportoletti et al., 2008). We therefore conclude that loss of *npm1a* would lead to increased genomic instability and acquisition of secondary mutation over time. Once HSPC clones acquire sufficient combinations of pathogenic mutations in hematopoietic genes, they are conferred a competitive advantage that permits their clonal expansion and would eventually trigger MDS/AML. The acquisition of secondary mutation may correlate with the appearance of disease noted at 6mpf in this study and would be interesting to follow up with further single cell multi-omics studies to investigate the pathways dysregulated downstream of specific mutation combinations. This will pave the way to undertake drug screening studies and develop more personalized treatments to help treat marrow failure and disease progression to MDS/AML.

## Materials and Methods

### Zebrafish

AB* zebrafish strains, along with transgenic strains and mutant strains, were kept in a 14/10 h light/dark cycle at 28°C. We used the following transgenic animals: *cd41*:GFP (Lin et al., 2005), *Tg(flk1:Hsa.HRAS-mCherry)^s896^* (Chi et al., 2008), *mpx*:GFP^c264Tg^ (Renshaw et al., 2006), *mpeg1:GFP^gl22^* (Ellett et al., 2011). We used the following mutant animals: *gata2a* (Dobrzycki et al., 2020). Experimental procedures in adult fish were done in accordance with the Animal Scientific Procedure Act 1986 under an approved Home Office Project License (PPL2470547).

### May-Grundwald (MG) and Giemsa staining WKM smears

Whole kidney marrow tissue was dissected from sacrificed animals and smeared on a superfrost plus slide (Thermo Scientific) and left to air dry for at least 1h at RT. MG (Sigma) was diluted 1:1 and Giemsa (Sigma) 1:10 with distilled water. Slides were incubated with diluted MG for 5 min at RT, then washed and repeated with diluted Giemsa for 30 min. Slides were washed well with distilled water and left to air dry before being mounted by DPX (Sigma) and imaged using a Leica DM750 scope with Leica ICC50W digital camera attachment.

### FACS

Cells were countered using a haemocytometer (Marionfield) and dead cells were stained using trypan blue (sigma) diluted 1:10. Cells were disassociated and re-suspended in 10% FCS (GIbco) 1xPBS and analysed using a BD LSRFORTESSA X-20 and dead cells were excluded using Hoechst’s (Invitrogen), used at 1:10000. Cells were sorted using BD FACSAria™ Fusion. For γH2AX, WKM cells were fixed in 4% PFA for 10 mins at RT, permibilised in 0.1% Triton for 5 mins at RT, blocked in 4% FBS for 20 mins at RT and stained with H2A.XS139ph (GeneTex, GTX127342) (diluted 1:1000 in 3% FCS/PBS) followed by AlexaFluro 594 goat anti-rabbit (Invitrogen, A11012), (diluted 1:1000 in 3% FCS/PBS).

### scRNA-Seq

3′-scRNAseq was completed using Chromium Next GEM single cell 3′ GEM library and Gel bead Kit v3.1 (10x Genomics) and sequenced using a NextSeq 500 (Illumina) at Genomics Birmingham (University of Birmingham). Analysis was completed using CellRanger, Loupe and R: Using Monocle 3 (Cao et al., 2019; Qiu et al., 2017a; Qiu et al., 2017b; Trapnell et al., 2014) and Seurat (Butler et al., 2018; Satija et al., 2015; Stuart et al., 2019). Results were presented using Graphpad Prism (v9). GO enrichment term analysis was completed biological function and using Metascape (Zhou et al., 2019).

### Immunofluorescence

WKM cells were cytospun onto superfrost plus slide (Thermo Scientific) and left to air dry for 15min at RT. Slides were pre-extracted (with pre-extraction buffer: 20mM NcCl, 3mM MgCl2, 300mM sucrose, 10mM PIPES, 0.5% Triton X-100) on ice for 5 min and then fixed in 4% PFA for 10min at RT. Slides were washed in PBS 3 times and then blocked in 100ul blocking buffer (10% FCS/PBS) for 1 hour at RT. H2A.XS139ph (GeneTex, GTX127342), 1:500 (diluted in 3% FCS/PBS), was added to the slide for 1 hour at RT, washed in PBS and then incubated with AlexaFluro 594 goat anti-rabbit (Invitrogen, A11012), 1:1000 (diluted in 3% FCS/PBS). Slides were then washed in PBS and mounted using DAPI mounting medium (abcam, ab104139).

### Cell culture and Western blot

HPC7 murine multipotent progenitor cells were obtained from Prof J. Frampton (University of Birmingham), and maintained in StemPro-34 SFM (Invitrogen, 10639011) containing supplement, 1% Pen/Strep, 1 x L-glutamine and 100 ng/ml SCF. Cells were treated with the NPM1 oligomerization inhibitor (Stratech Scientific Ltd) (Qi et al., 2008) (1 uM for 24 h), harvested by centrifugation and whole cell extracts were obtained by lysis in UTB buffer (8 M Urea, 50 mM Tris, 150 mM β-mercaptoethanol). Lysates were separated and analysed by SDS-PAGE following standard procedures. The following antibodies were used: γH2AX (Millipore; cat#05-636; RRID:AB_309864), H2A (Millipore; cat#07-146; RRID:AB_11212920), Anti-rabbit HRP (Agilent; cat#P0399; RRID: AB_2617141), Anti-mouse HRP (Agilent; Cat# P0447; RRID: AB_2617137).

## Acknowledgements

We thank the staff of the Biomedical Services Unit for fish husbandry. We thank Dr Adriana Flores-Langarica and Dr Paola Pietroni from the flow cytometry facility (University of Birmingham) for cell sorting. We thank the staff of Genomics Birmingham (University of Birmingham) for genomic sequencing and Prof. J. Frampton for the HPC7 cells. This research was supported by a Welcome Trust ISSF Award to R.M. and by the British Heart Foundation (BHF IBSR Fellowship FS/13/50/30436 to R.M. and C.B.M.).

