## Supplemental Figures for "Gata2a maintains *cebpa* and *npm1a* in haematopoietic stem cells to sustain lineage differentiation and genome stability"

### Figure S1

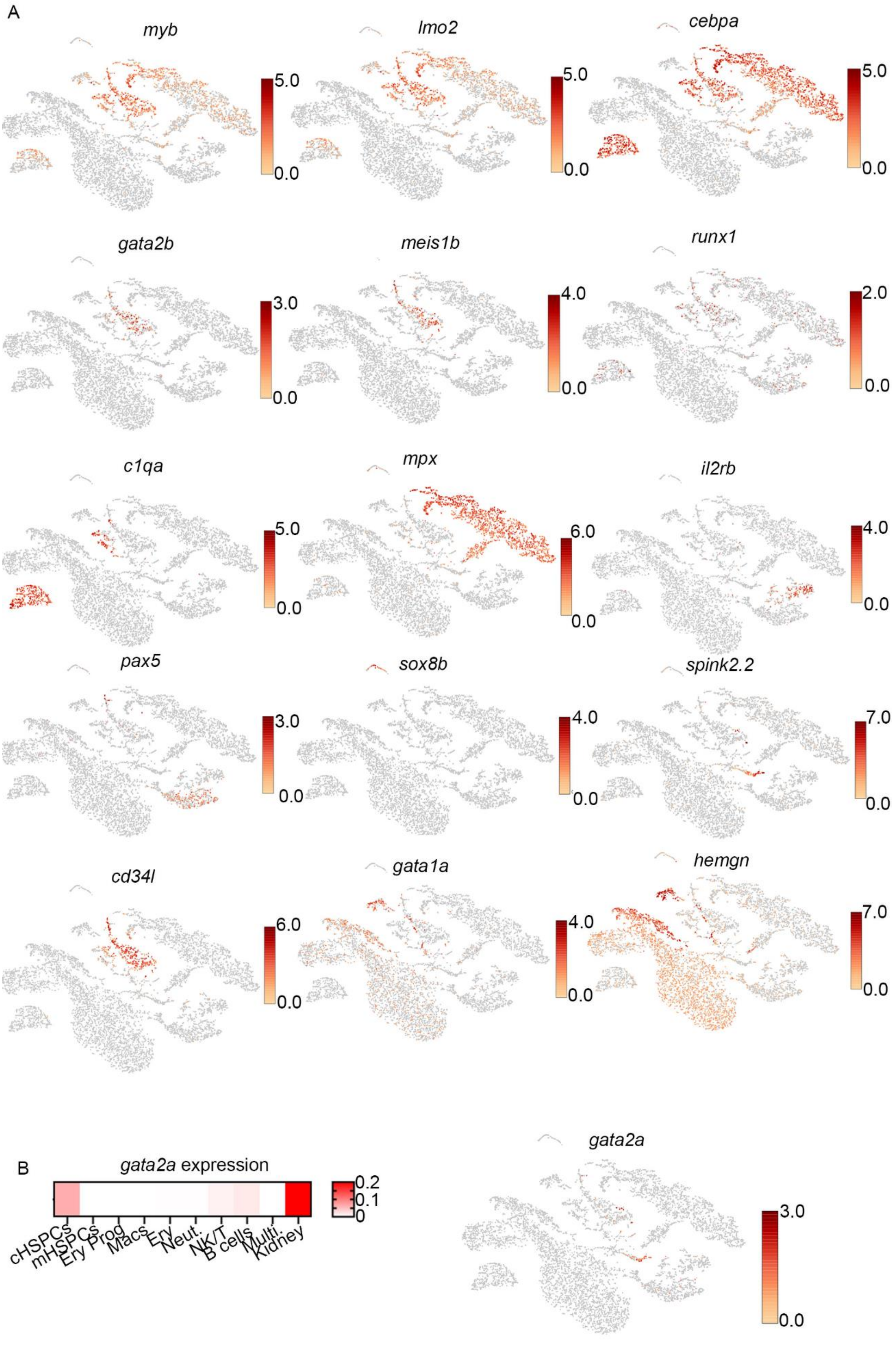

A
myb
5.0
0.0
lmo2
5.0
0.0
cebpa
5.0
0.0
gata2b
3.0
0.0
meis1b
4.0
0.0
runx1
2.0
0.0
c1qa
5.0
0.0
mpx
6.0
0.0
il2rb
4.0
0.0
pax5
3.0
0.0
sox8b
4.0
0.0
spink2.2
7.0
0.0
cd34l
6.0
0.0
gata1a
4.0
0.0
hemgn
7.0
0.0
B
gata2a expression
0.2
0.1
0
cHSPCs
mHSPCs
Ery Prog
Macs
Ery
Neut.
NK/T
B cells
Multi.
Kidney
gata2a
3.0
0.0

**Figure S1. Identification of haematopoietic populations in WKM.**
(A) tSNE clustering of WKM showing expression levels of indicated genes. Expression levels are log2 fold expression per cell (each point represents one cell). (B) Heat map of mean log2 fold *gata2a* expression per indicated cluster and tSNE clustering of WKM showing expression levels of *gata2a* (expression levels are log2 fold expression per cell (each point represents one cell)).

**Figure S2**

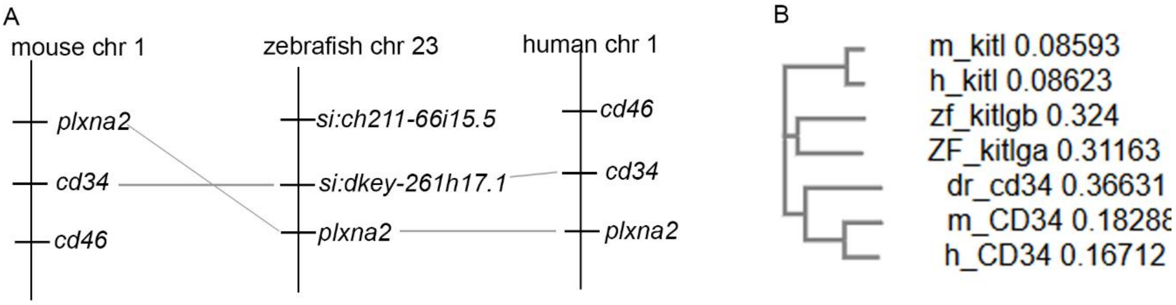


C

```
dr_cd34    MAVR-------RMNELWKSMPLALMFGLLLLHGCQGQDEAGASTTSQ-SVGAE----TNP    48
m_Cd34     MQVHRDTRAGLLLPWRWVALCLMSLLHL----NNL-----TSATTETSTQGISPSVPTNE    51
h_CD34     MLVRRGARAGPRMPRGWTALCLLSLLPS----GFMSLDNNGTATPELPTQGTFSNVSTNV    56
           * *:        :   * :: *  ::      .        ::* .  : *      **

dr_cd34    TTAAAMIADVQGT----------EQPATSGLETQSNYTLS--------------------    78
m_Cd34     SVEENITSSIPGSTSHYLIYQDSSKTTPAISETMVNFTVTSGIPSGSGTPHTFSQPQTSP    111
h_CD34     SYQETTTPSTLGSTSLHPVSQHGNEATTNITETTVKFTSTSVITSVYGNTNSSVQSQTSV    116
           :       .  *:          .: :    **  ::* :

dr_cd34    -GTAVTNQE---IIEKTTNSSLMP--------QQGN------INPTTVN--PTPE-RKNN    117
m_Cd34     TGILPTTSDSISTSEMTWKSSLPSINVSDYSPNNSSFEMTSPTEPYAYTSSSAPSAIKGE    171
h_CD34     ISTVFTTPANVSTPETTLKPSLSPGNVSDLSTTSTSL-ATSPTKPYTSS-SPILSDIKAE    174
            .   *.       * * : **           . .       :* : .     .  * :

dr_cd34    I----TDDSKGGRGVISDQSETSINETVK-------------------------------    142
m_Cd34     IKCSGIREVRLAQGICLELSEASSCEEFKKEKGEDLIQILCEKEEAEADAGASVCSLLLA    231
h_CD34     IKCSGIREVKLTQGICLEQNKTSSCAEFKKDRGEGLARVLCGEEQADADAGAQVCSLLLA    234
           *      : :  :*:  : .::*    .*

dr_cd34    ---------VIN-------------FDGKQETSKESGSSDAKAKADKDTTS-GGSSSAVF    179
m_Cd34     QSEVRPECLLMVLANSTELPSKLQLMEKHQSDLRKLGIQSF---NKQDIGSHQSYSRKTL    288
h_CD34     QSEVRPQCLLLVLANRTEISSKLQLMKKHQSDLKKLGILDF---TEQDVASHQSYSQKTL    291
                    ::              :. :*.  :: *  .     .:*  *  . *  .:

dr_cd34    VSVLLTGLLLAGIIVGIYYFKCHRRTNGKGMKLAEESYTADEENQGNTLVSVAPLNQPEP    239
m_Cd34     IALVTSGVLLAILGTTGYFLMNRRSWSPTGERLGEDPYYTE--NGGGQGYSSGPGASPET    346
h_CD34     IALVTSGALLAVLGITGYFLMNRRSWSPTGERLGEDPYYTE--NGGGQGYSSGPGTSPEA    349
           :::: :* *** :    *::  :*  . .* :*.*: * ::  * *.   * .*  .**

dr_cd34    QEKPSLNGESQEAVKPQTPPAATNGHSTT--KTADTEL      275
m_Cd34     QGKANVTRGAQENGTGQAT--SRNGHSARQHVVADTEL      382 25.87% compared to dr
h_CD34     QGKASVNRGAQENGTGQAT--SRNGHSARQHVVADTEL      385 28.79% compared to dr
           * * .:.  :**  . *:   : ****:    .*****
```

Underlined in red: CD34 antigen- as defined by ensembl

**Figure S2. Identification of *cd34l* in zebrafish.**
(A) Synteny analysis of cd34l/ *si:dkey-261h17.1* across human, mouse and zebrafish. (B) Phylogentic analysis of *cd34 and kitl* across human, mouse and zebrafish. (D) Alignment of human, mouse, and zebrafish cd34l/ *si:dkey-261h17.1* with Clustal Omega (EMBL-European Bioinformatics Institute). Predicted CD34 antigen underlined in red (ENSEMBL). Zebrafish *cd34l* shows 25.87% amino acid similarity with mouse *CD34* and 28.79% amino acid similarity with human *CD34* (analysed using Clustal Omega). Chr: chromosome, dr: *danio rerio*,

### Figure S3

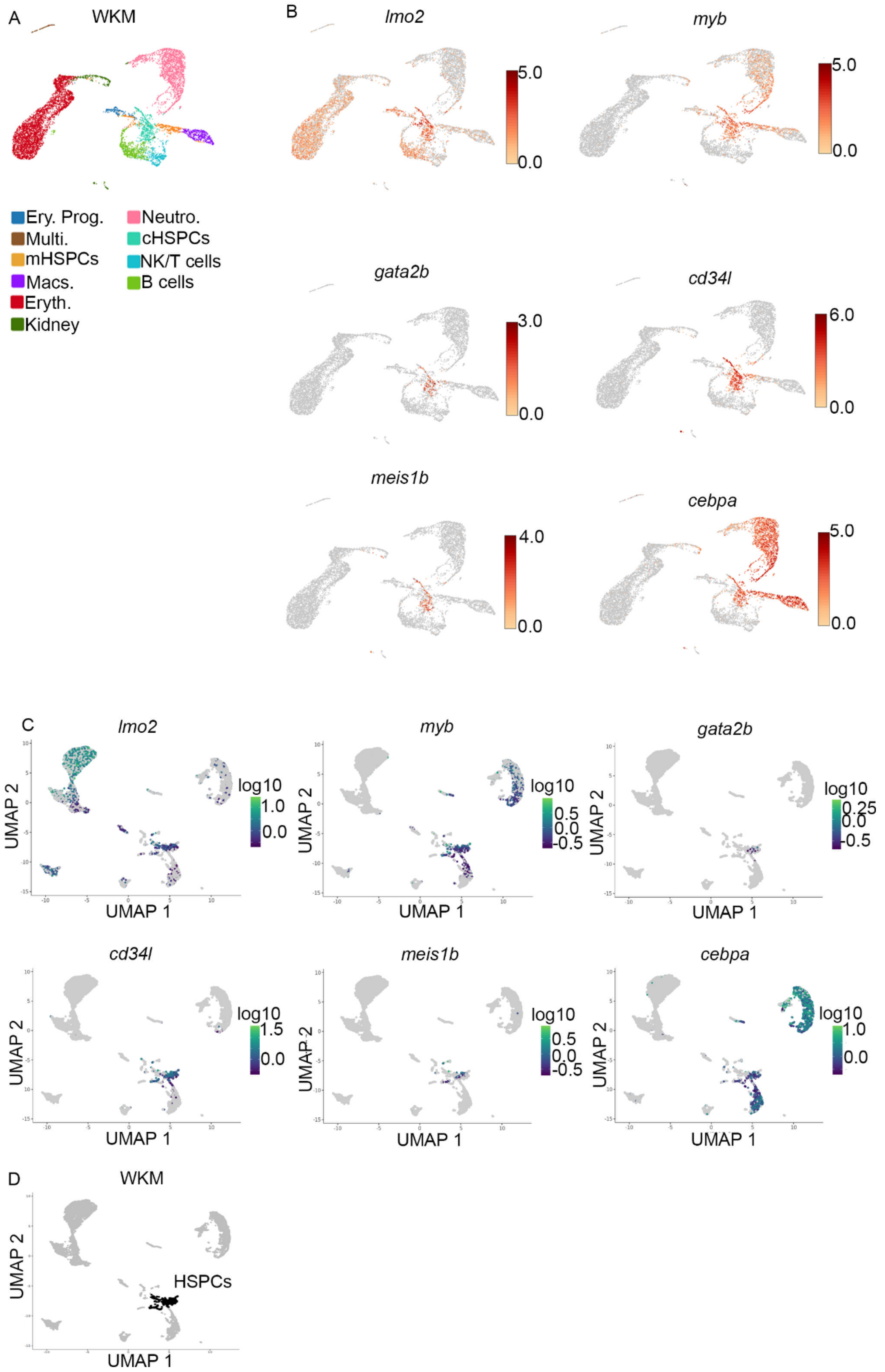
A
WKM
Ery. Prog.
Multi.
mHSPCs
Macs.
Eryth.
Kidney
Neutro.
cHSPCs
NK/T cells
B cells
B
lmo2
myb
gata2b
cd34l
meis1b
cebpa
C
D
HSPCs
UMAP 1
UMAP 2
log10

**Figure S3. Selecting HSPCs for pseudotime analysis.**

(A) UMAP plot of WT and *gata2a*$^{-/-}$ WKM cells at 5mpf (data from fig. 2A) indicating the main haematopoietic populations. (B) UMAP (using Loupe browser) clustering and identification of HSPCs based on the expression of *lmo2*, *myb, gata2b, cd34l, meis1b* and *cebpa*. Expression levels are log2 fold expression per cell (each point represents one cell). UMAP (using Loupe browser) clustering and identification of HSPCs based on the expression of *lmo2*, *myb, gata2b, cd34l, meis1b* and *cebpa*. Expression levels are log2 fold expression per cell (each point represents one cell). (C) UMAP (using Monocle 3) clustering and identification of HSPCs based on the expression of *lmo2*, *myb, gata2b, cd34l, meis1b* and *cebpa*. Expression levels are log10 fold expression per cell (each point represents one cell). (D) Selection of HSPCs in monocle 3.

**Figure S4**

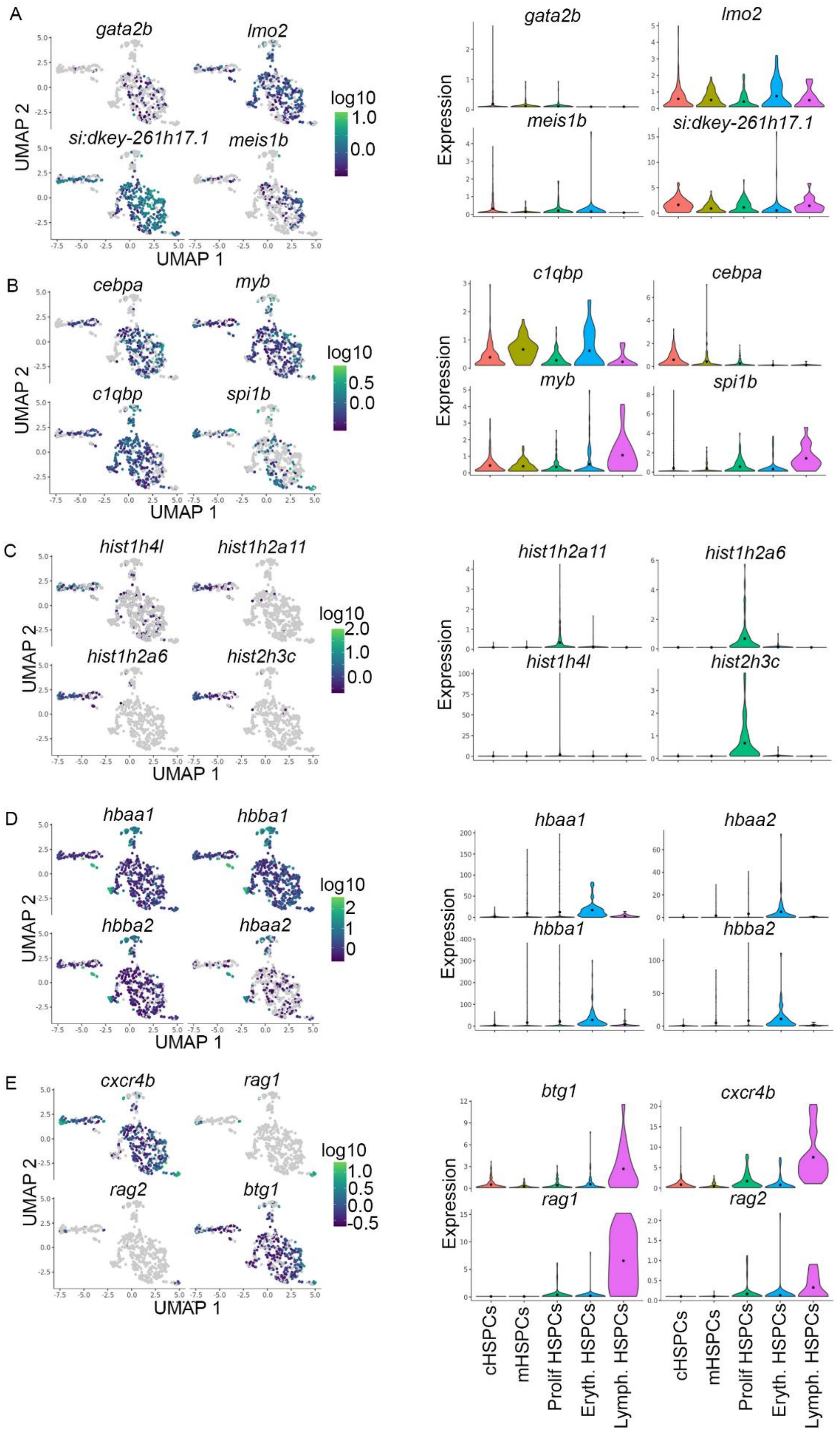
A
gata2b
lmo2
si:dkey-261h17.1
meis1b
log10
UMAP 2
UMAP 1
Expression
B
cebpa
myb
c1qbp
spi1b
C
hist1h4l
hist1h2a11
hist1h2a6
hist2h3c
D
hbaa1
hbba1
hbba2
hbaa2
E
cxcr4b
rag1
rag2
btg1
cHSPCs
mHSPCs
Prolif HSPCs
Eryth. HSPCs
Lymph. HSPCs

**Figure S4. Characterising sub populations of HSPCs in Monocle 3.**
(A-E) UMAP clustering and violin plots (using Monocle 3) to identify HSPC sub populations based on the expression of indicated markers. Expression levels are log10 fold expression per cell. cHSPCs: core HSPCs, mHSPCs: myeloid primed HSPCs, Prolif. HSPCs: proliferating HSPCs, Eryth. HSPCs: erythroid primed HSPCs, Lymph. HSPCs: lymphoid primed HSPCs.

**Figure S5**

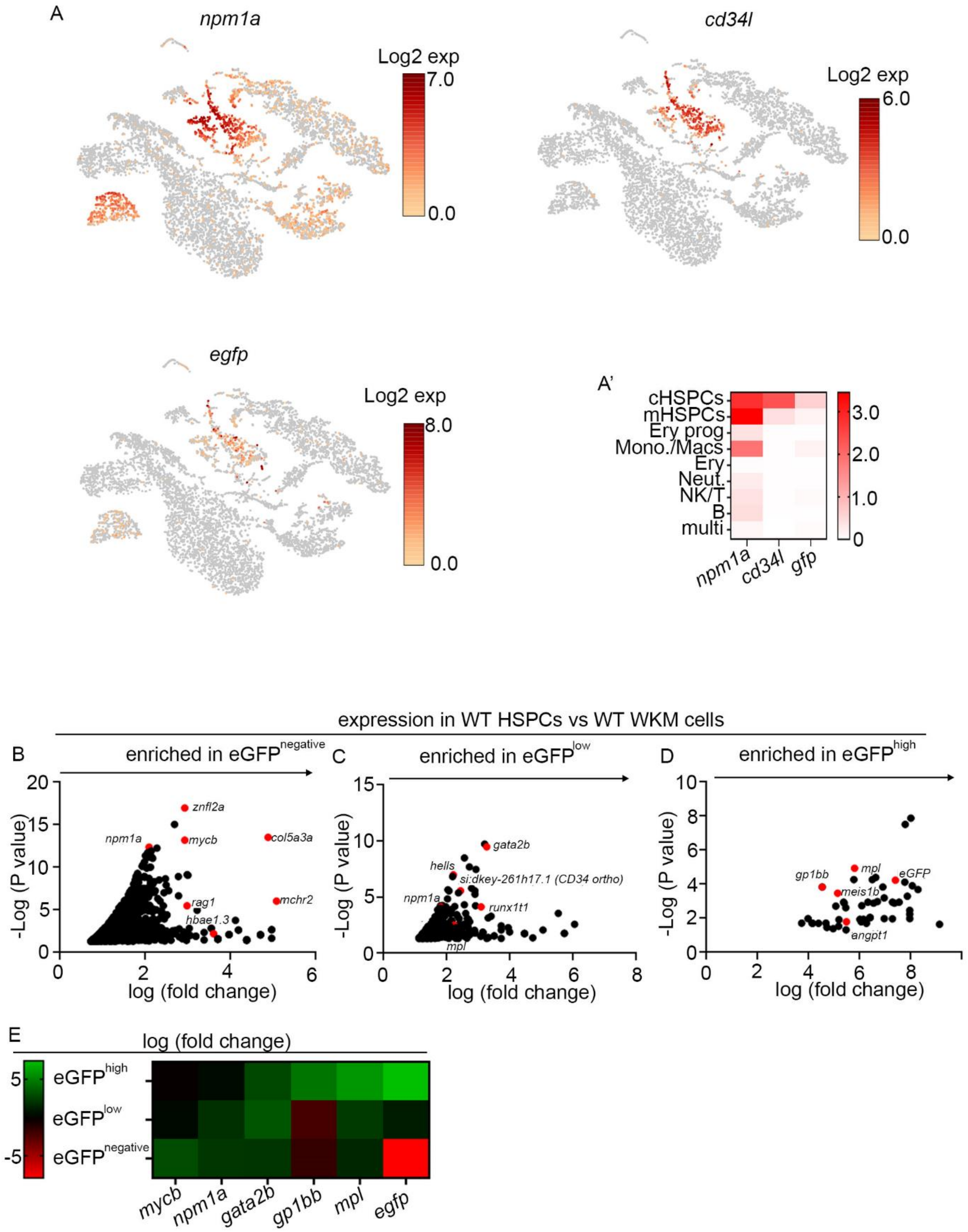


**Figure S5. Cd41:GFP$^{low}$ cells are enriched in haematopoietic stem cell markers.**
(A) tSNE clustering of WKM showing expression levels of indicated genes. Expression levels are log2 fold expression per cell (each point represents one cell). (A') Heat map showing mean log2 fold expression levels of genes per indicated cluster. (B-D) Volcano plots showing log2 fold change vs –log10 (p value, determined by using the Benjamini-

Hochberg correction for multiple tests) in GFP$^{negative}$, GFP$^{low}$, GFP$^{high}$. Each dot represent one gene, only genes with a p value < 0.05 are shown. (E) heatmap showing mean expression levels of selected genes in GFP populations. Expression if log2 fold change.

**Figure S6**

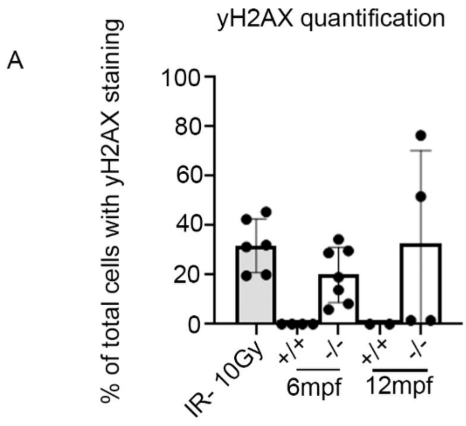


**Figure S6. γH2AX staining is increased in *gata2a*$^{-/-}$ WKM cells.**
(A) Percentage of total cells within the imaged region that contains γH2AX staining.
